# Engineering RNA viruses with unnatural amino acid to evoke adjustable immune response in mice

**DOI:** 10.1101/2021.12.04.471206

**Authors:** Zhetao Zheng, Yu Wang, Xuesheng Wu, Haoran Zhang, Hongmin Chen, Haishuang Lin, Yuxuan Shen, Qing Xia

## Abstract

Ribonucleic acid (RNA) viruses pose heavy burdens on public-health systems. Synthetic biology holds great potential for artificially controlling their replication, a strategy that could be used to attenuate infectious viruses but is still in the exploratory stage. Herein, we used the genetic-code expansion technique to convert *Enterovirus* 71 (EV71), a model of RNA virus, into a controllable EV71 strain carrying the unnatural amino acid (UAA) Nε-2-azidoethyloxycarbonyl-L-lysine (NAEK), which we termed an EV71-NAEK virus. EV71-NAEK could recapitulate an authentic NAEK time- and dose-dependent infection *in vitro* and *in vivo*, which could serve as a novel method to manipulate virulent viruses in conventional laboratories. We further validated the prophylactic effect of EV71-NAEK in two mouse models. In susceptible parent mice, vaccination with EV71-NAEK elicited a strong immune response and potentially protected their neonatal offspring from lethal challenge similar to that of commercial vaccines. Meanwhile, in transgenic mice harboring a PylRS-tRNA^Pyl^ pair, substantial elements of genetic-code expansion technology, EV71-NAEK evoked an adjustable neutralizing-antibody response in a strictly external NAEK dose-dependent manner. These findings suggested that EV71-NAEK could be the basis of a feasible immunization program for populations with different levels of immunity. Moreover, we expanded the strategy to generate controllable coxsackieviruses and severe acute respiratory syndrome coronavirus 2 (SARS-CoV-2) for conceptual verification. In combination, these results could underlie a competent strategy for attenuating viruses and priming the immune system via artificial control, which might be a promising direction for the development of amenable vaccine candidates and be broadly applied to other RNA viruses.

## Introduction

RNA viruses account for the majority of human viral infections^1^ and remain difficult to control despite the availability of several working vaccine strategies^2–4^. While synthetic biology holds great potential for biomedical engineering, it is limited in accessing vector design for gene therapy^5–8^, which renders it far from practical in vaccine production^9, 10^. Therefore, by integrating advances in reverse genetics^11, 12^ with insights from system vaccinology^13, 14^, viral-engineering methods could be adapted to effective and smart vaccine design in a novel way.

Furthermore, a replication-incompetent virus that holds artificial amber codons in its genome and propagates using genetic-code expansion technology^15, 16^ elicits robust immunity in animal models and subverts traditional vaccine approaches^17, 18^. Nevertheless, the decisive step is the incorporation of unnatural amino acid (UAA) into the nascent viral polypeptide at the desired positions, which are selected by massive random mutations with undefined rules^18^, impeding the approach expanded to other emerging viruses. Moreover, UAA modification is a switch that enables artificial control of virus replication^19, 20^, hinting at a potential strategy of precisely controlling viral replication *in vivo* to imprint desired antibody landscapes.

We applied the genetic-code expansion technique to modify *Enterovirus* 71 (EV71) as a model to (a) collect data that would define rules for UAA incorporation and (b) test whether the optimized EV71-UAA would require the conservation of the full infectious form for immunogenicity and replication under artificial control *in vitro* and *in vivo* for safety^21, 22^. We further exploited the virus to evoke strong and adjustable antibody responses in normal and transgenic mice harboring PylRS-tRNA^Pyl^ pairs as a potential approach to vaccinating susceptible individuals with differing levels of immunity^23^.

## Results

### *In vitro* UAA-controllable RNA virus-packaging and -production system

In this study, we chose EV71, the major pathogen in hand-foot-mouth disease (HFMD) in children and infants^24, 25^, as a model virus to test the feasibility of a production pipeline for viruses with expanded genetic codes. An amber codon artificially introduced into the viral genome renders the virus replication incompetent in conventional cells but recovers replicative potential using UAA incorporation machinery, which is composed of three biorthogonal elements: aminoacyl-tRNA synthetases (aaRS), transfer RNA (tRNA), and UAA **(Fig. 1a)**. To improve the compatibility of the incorporation machinery with production of RNA viruses carrying UAA, we designed a two-round viral-production pipeline: (a) viral RNA into which an amber codon was introduced was transfected into human embryonic kidney 293T (HEK293T) cells integrated with well-established orthogonal MmPylRS/tRNA^MmPyl^ pairs (NAEK-HEK293T cells) for first-round viral packaging **(Fig. 1a, left)**; (b) propagation of a NAEK-Vero stable cell line for high-yield viral production in the presence of NAEK **(Fig. 1a, right)**. Given that the NAEK-HEK293T stable cell line was previously reported^18^, we first sought to establish the NAEK-Vero line by applying the *Desulfobacula toluolica* (Tol2) transposon to integrate the orthogonal pair into the Vero-E6 cells genome **(Extended Data Fig. 1)**. We confirmed the successful expression of MmPylRS, tRNA^Pyl^, and NAEK-dependent GFP^39TAG^ carrying amber-codon replacements in 39 sites, facilitating selection of cell lines with efficient UAA translation machinery **(Fig. 1b, c)**. In addition, we observed that the machinery’s introduction did not significantly affect cell proliferation in either line, which was beneficial for viral packaging and propagation **(Fig. 1d)**.

**Figure 1.**
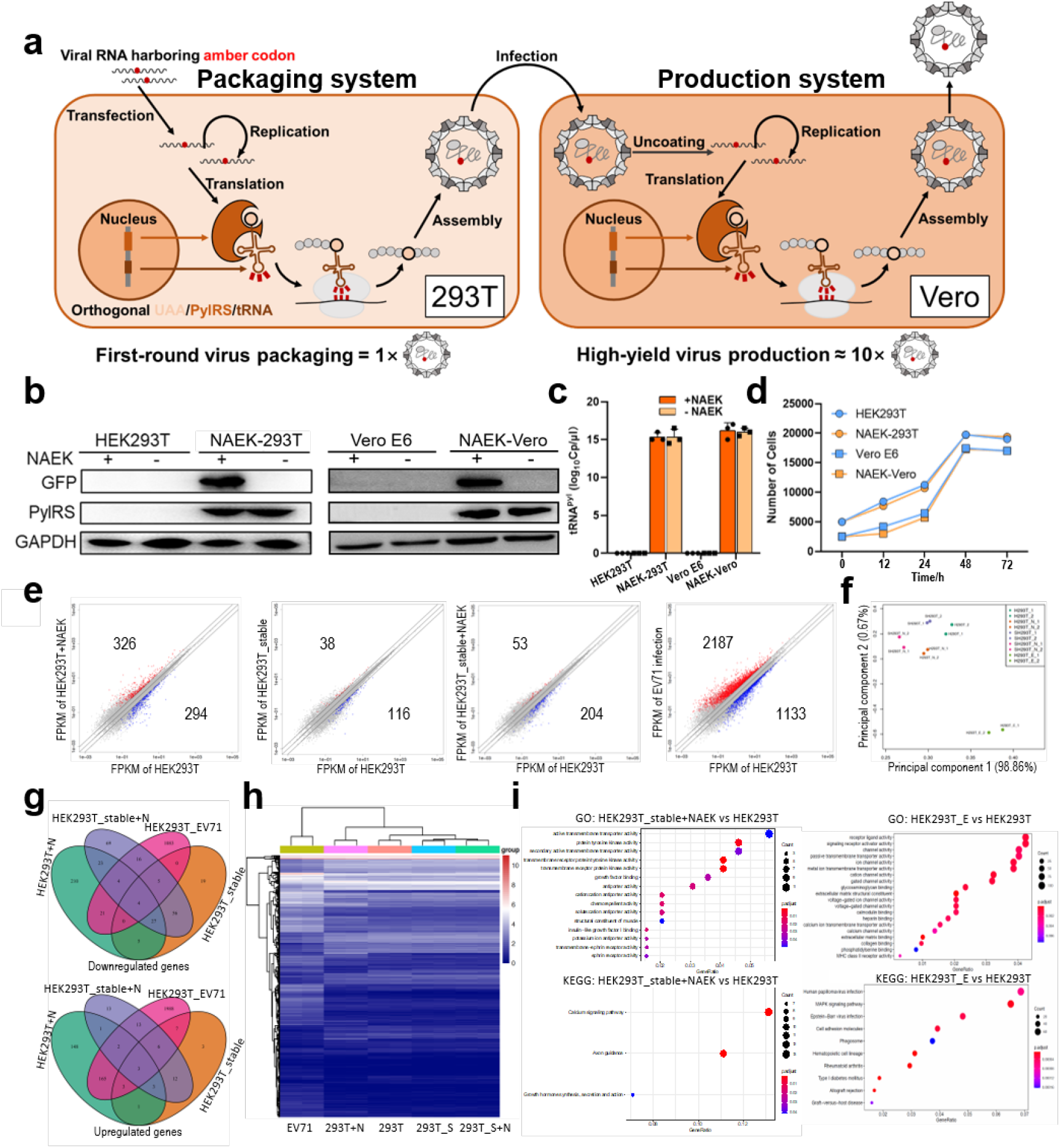
Construction and identification of UAA packaging system to generate EV71-NAEK virus. (**a**) Schematic illustration of EV71-NAEK viral packaging and production. An artificial amber codon was introduced into the viral genome to create a replication-incompetent virus using engineered HEK293T cells harboring a UAA incorporation system composed of orthogonal PylRS, tRNA^Pyl^, and the relevant UAA (NAEK in this study). The UAG was read-through by the UAA incorporation system, generating EV71-NAEK virus, which was then produced on a large scale by Vero-E6 cells harboring the UAA system. (**b**) The characterization of UAA packaging system by WB analysis. (**c**) Reverse transcription polymerase chain reaction (RT-PCR) validation of the tRNA^Pyl^ transcripts. Data are represented as mean ± standard deviation (SD; n = 3). (**d**) The effect of engineered HEK293T and Vero-E6 cells harboring the UAA incorporation system on cell proliferation. (**e**) WTA of HEK293T cells, the engineered packaging systems (HEK293T_stable), and EV71-infected HEK293T cells. The plots show whole-transcriptome fragments per kilobase of exon per million fragments mapped (FPKM). Red dots indicate upregulated genes; blue dots downregulated genes. Two biological replicates were used per sample. (**f**) PCA of HEK293T±NAEK, HEK293T_stable (SHEK293T) ±NAEK and EV71-infected HEK293T cells (HEK293T_E). (**g**) Venn diagram showing significant overlap (*P* < 0.005) of DEGs among HEK293T cells, engineered packaging systems, and EV71-infected HEK293T cells. (**h**) Hierarchical-clustering and heatmap analysis of DEGs in HEK293T cells, the engineered packaging systems, and EV71-infected HEK293T cells. (**i**) Representative GO terms and KEGG enrichment analysis of the engineered HEK293T+NAEK and EV71-infected HEK293T cells compared with HEK293T cells.

After establishing appropriate viral-packaging systems in the stable cell lines, we evaluated the effect of the NAEK system’s incorporation on gene expression via whole-transcriptome analysis (WTA). Whole-transcriptome plots and principal-component analysis (PCA) revealed that the cell line with engineered systems had gene expression patterns similar to those of HEK293T cells, which differed significantly from those of EV71-infected HEK293T cells (**Fig. 1e, f**). The Venn diagram (**Fig. 1g**) shows that the number of differentially expressed genes (DEGs) in EV71-infected HEK293T cells was significantly higher than that in the stable cell line ± NAEK groups. Hierarchical-clustering analysis of DEGs showed two main clusters, which are displayed in the heatmap, further indicating that the stable cell line ± NAEK groups had DEGs similar to those of HEK293T cells, which in turn were very different from those of EV71- infected HEK293T cells (**Fig. 1h**). Gene Ontology (GO) terms and Kyoto Encyclopedia of Genes and Genomes (KEGG) signaling pathways further indicated their common application of stable cell line ± NAEK groups for generating artificial viruses from other extant RNA viruses (**Fig. 1i, Extended Data Fig. 2a, b**). The WTA results were similar in the NAEK-Vero stable cell line (**Extended Data Fig. 3**). The UAA system displayed the same DEG expression patterns, GO terms, and KEGG signaling pathways as in HEK293T cells, indicating that the effect of UAA incorporation was far less than that of viral infection and that the system was appropriate for controllable viral production.

### Parallel screening revealed rules for NAEK incorporation

A guideline for identifying optimal amber-codon introduction into the viral genome based on reliable NAEK incorporation machinery is pivotal to an effective NAEK- controllable viral-replication system. Therefore, we profiled the entire viral genome by generating different EV71-NAEK viruses containing mutant amber codons and observed the varying viral titers **(Fig. 2a)**. Although all encoding genes except 2B contributed to at least one mutant codon (2 of 10 codons in VP4, 1 of 4 in VP2, 1 of 4 in VP3, 5 of 10 in VP1, 1 of 5 in 2A, 0 of 3 in 2B, 3 of 10 in 2C, 1 of 4 in 3A, 1 of 3 in 3B, 5 of 5 in 3C, and 9 of 10 in 3D) whose NAEK-controllable virus could be successfully packaged and propagated, we noted that viruses in 3C or 3D genes showed prevalently higher viral titers **(Fig. 2a, Extended Data Fig. 4)**. To assess the features of incorporated positions that could affect viral titer, we further tested 15 and 20 mutant codons located in particular EV71-3C and 3D genes, respectively **(Fig. 2b, c)**. We thus generated 9 and 11 variants, respectively, with virions of 10^4^–10^6.5^ TCID50/mL, and clearly observed NAEK-dependent cytopathic-effect (CPE) phenotypes as expected (Fig. 2d, Extended Data Fig. 5).

**Figure 2.**
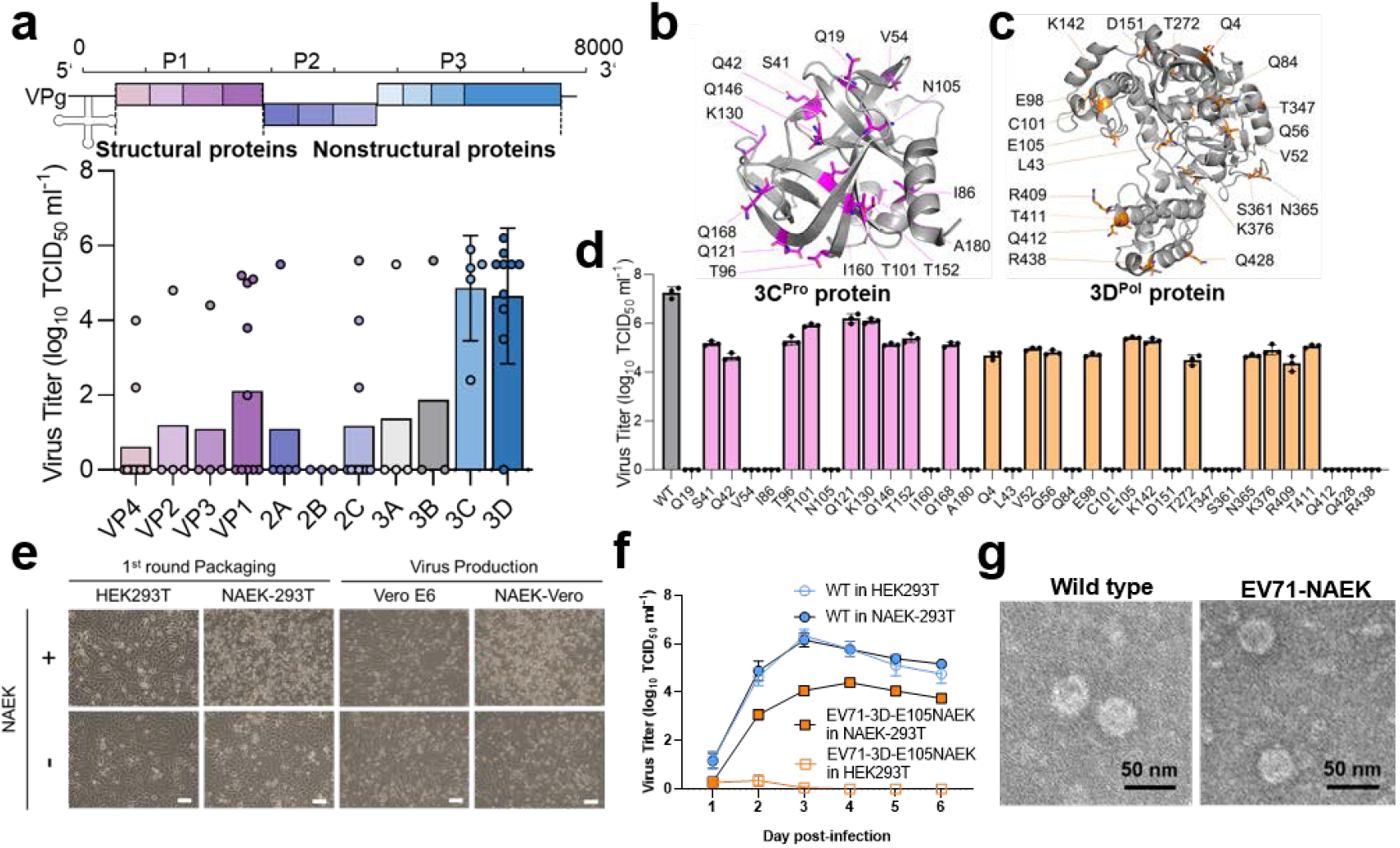
Screening for optimal incorporation sites. (**a**) Optimal amber insertion genes (including VP1–VP4, 2A–2C, and 3A–3D proteins) were discovered by profiling the viral genome, which turned out to be the gene for 3C/3D protein. (**b–d**) We tested 35 locations in the gene of variable, average, and conserved domains in EV71-3C/3D protein and identified the optimal mutant site for the UAA system to read through. That site was 3D-E105NAEK, due to its relevant high packaging efficiency and viral titer. (**e**) CPE of EV71-3D-E105NAEK packaged with the HEK293T-NAEK system and produced by the VeroE6-NAEK system in the presence or absence of NAEK. Scale bars, 50 μm. (**f**) Growth kinetic curves of EV71-3D-E105NAEK generated by the NAEK system in the presence or absence of NAEK at indicated time points. Data are represented as mean ± SD (n = 3). (**g**) Morphologies of the EV71-3D-E105NAEK and WT EV71 viruses as shown by TEM. Scale bars, 50 nm.

These results indicated that incorporating UAA to replace amino acids with similar properties (NAEK to lysine or glycine) and appropriate conservation could increase the production efficiency and genomic fidelity of controllable viruses (**Extended Data Fig. 6a–c**). Of the 35 incorporation positions screened in this study, 30 residues were located on the surface of 3C/3D, and the other five were inaccessible to the solvent (**Extended Data Fig. 7**). Furthermore, surface sites might be more open to NAEK with respect to amino acid similarity (**Extended Data Fig. 6a–c**). Other factors, such as residue– residue interaction or steric effects, also affected overall package efficiency to some extent. Therefore, we performed a logistic-regression analysis (**Extended Data Fig. 6d–f**), the results of which indicated that viral packaging correlated positively and significantly (W4 > 10, *P* < 0.05) with amino acid similarity. Therefore, when the original amino acid at a certain position resembled NAEK, the respective mutant amber codon tended to be better read-through by the corresponding system, rendering controllable viral production more possible. In addition, the NAEK system was also sensitive to the conservation level (W3 = −3.6, *P* < 0.05), indicating that NAEK could be better incorporated into the less conserved site of a viral protein.

### Prediction and validation of UAA-controllable virus design

We tested whether the aforementioned logistic function could be applied to other RNA viruses to guide mutant amber-codon selection. Owing to the coronavirus 2019 (COVID-19) pandemic, we considered severe acute respiratory syndrome coronavirus 2 (SARS-CoV-2) as an ideal test virus (**Extended Data Fig. 8**). Because we were unqualified to handle this Biosafety Level 3 (BSL-3) virus, we tentatively introduced amber codons into its synthesized nucleocapsid (N) genes, including N29^TAG^, G97^TAG^, E136^TAG^, and A211^TAG^ (**Extended Data Fig. 8a**). As such, a fluorescent imaging and Western blot (WB) analysis confirmed that all these N proteins were successfully expressed in the stably transgenic cells to which NAEK had been administered (**Extended Data Fig. 8b, c**). The main types of *Enterovirus* that account for human infections, including the enteroviruses EV71 and EVD68 and the coxsackieviruses CA6, CA10, and CA16, comprised the P1, P2, and P3 genomes, respectively encoding the 3AB, 3C, and 3D proteins (**Extended Data Fig. 8d**). We thereby managed to expand our strategy to the abovementioned RNA viruses, demonstrating the feasibility of logistic function. We found similar NAEK-dependent CPE formation and viral RNA copies of those viruses compared with wild-type (WT) viruses (**Extended Data Fig. 8e–h**).

For EV71, we selected EV71-3D-E105NAEK, with one codon encoding Glycine in the 3D^P^°^l^ protein, for amber-codon replacement (3D-E105^TAG^), and attempted to survey its viral characteristics. After EV71-3D-E105NAEK infection of both cell lines, we observed the CPE phenotype, indicating that successful production and widespread propagation of EV71-3D-E105NAEK was NAEK dependent (**Fig. 2e**). Compared with that of WT EV71, the growth curve of EV71-3D-E105NAEK further demonstrated that NAEK-controllable viral replication was well established *in vitro* (**Fig. 2f**). Moreover, transmission electron microscopy (TEM) showed no significant differences in morphologies, implying that EV71-3D-E105NAEK maintained the original phenotype of parental EV71 (**Fig. 2g**). Owing to its combined high viral titer and low escape frequency, we chose EV71-3D-E105NAEK as a model strain for further study *in vivo*.

### Efficacy, safety, and protective evaluation of EV71-3D-E105NAEK as a vaccine candidate *in vivo*

Next, we evaluated the *in vivo* safety and immunogenicity of intraperitoneally (i.p.) injected EV71-3D-E105NAEK in adult BALB/c mice, using commercial Sinovic- EV71 vaccine as positive control (**Fig. 3a**). No adult mice died from EV71-3D- E105NAEK or showed obvious body weight (BW) loss or other health issues due to this injection (**Fig. 3b, c**). We detected no significant differences in viral load in the brain or skeletal muscle between the EV71-3D-E105NAEK and Sinovic-EV71 groups, and even less viral load in small intestine was observed in EV71-3D-E105NAEK mice than in Sinovic-EV71 mice, indicating the good *in vivo* safety of EV71-3D-E105NAEK as a vaccine candidate (**Fig. 3d**). Two weeks after single- or double-dose immunization, both the EV71-3D-E105NAEK and Sinovic-EV71 vaccines induced robust serum immunoglobulin M (IgM) and anti–viral protein 1 (VP-1) IgG (**Fig. 3e, f**). To determine the potential protective efficacy of EV71-3D-E105NAEK in neonatal mice, we injected a lethal EV71 strain (SD059) i.p. into the 4-day-old offspring of immunized BALB/c mice. All of these neonatal mice—in contrast with those in the vehicle group that succumbed within 14 days—survived after immunization with EV71-3D-E105NAEK and Sinovic-EV71 vaccines (**Fig. 3g**), with no significant BW loss (**Fig. 3h**). In addition, viral load in the small intestine, brain, and spinal cord in both the EV71-3D-E105NAEK and Sinovic-EV71 groups was significantly lower than in the vehicle group (**Fig. 3i**), indicating the protective efficacy of EV71-3D-E105NAEK in neonatal mice. Additionally, immunostaining and hematoxylin and eosin (H&E) staining of the small intestine showed no EV71 viral replication or pathological changes in the neonatal offspring of immunized BALB/c mice compared with the vehicle group (**Fig. 3j, k**). In brief, our method could produce a reliable EV71-3D-E105NAEK vaccine candidate, comparable to the commercial vaccine, in a mouse model.

**Figure 3.**
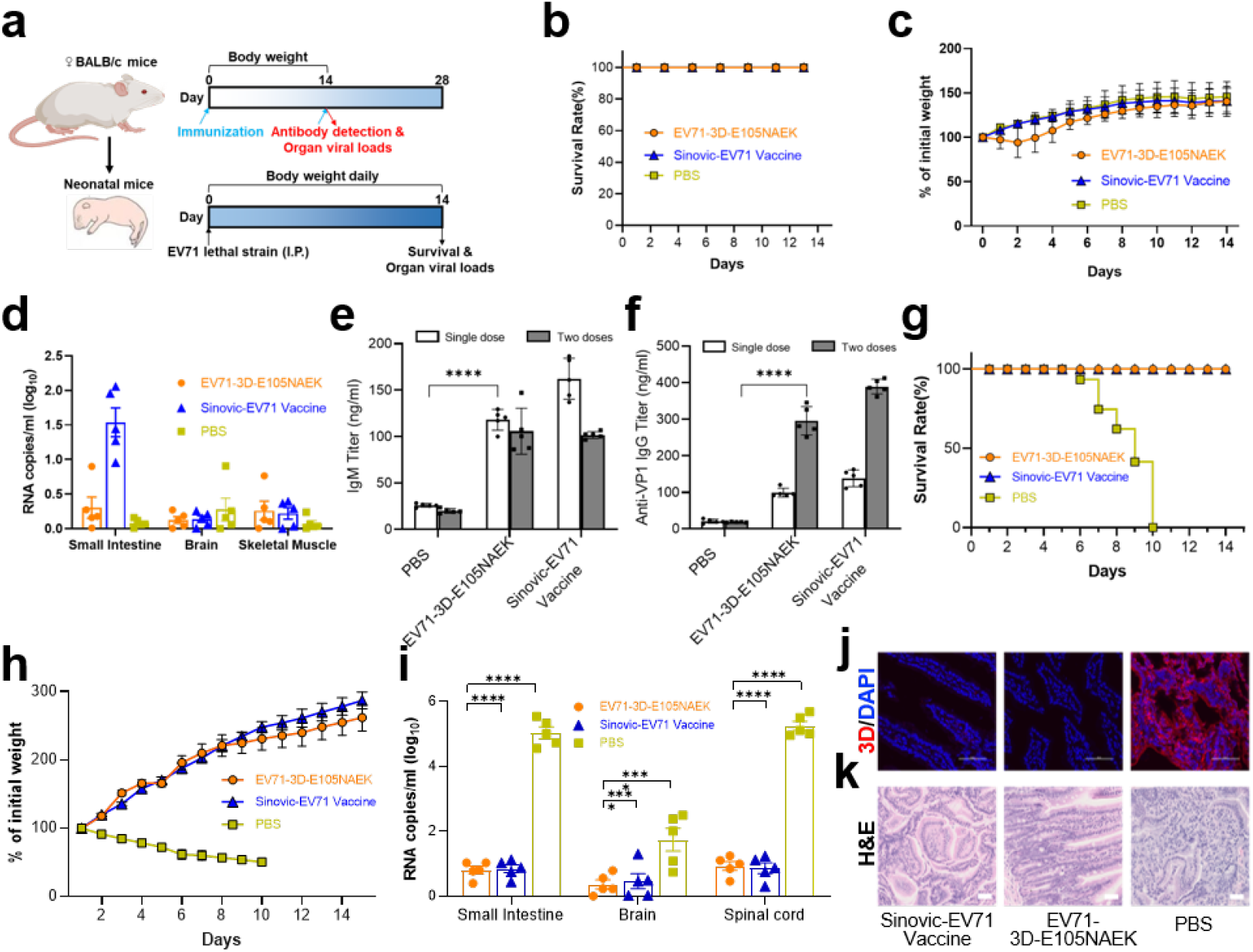
Evaluation of the EV71-3D-E105NAEK virus’s safety, immunogenicity, and protection levels. (**a**) Schematic of study design. To evaluate safety and immunogenicity, 4-week-old female BALB/c mice were first injected i.p. with EV71- 3D-E105NAEK for 14 days. To evaluate protection, the 4-day-old offspring of immunized BALB/c mice received i.p. injections of a lethal EV71 strain (SD059) for 14 days. (**b, c**) Survival rates and BW changes in BALB/c mice. (**d**) Viral RNA copies detected in the small intestine, brain, and skeletal muscle. (**e, f**) IgG and IgM titers detected before and after one or two doses of immunization. (**g, h**) Survival rates and BW changes in neonatal child BALB/c mice. (**i**) Detection of viral RNA copies in neonatal-mouse tissues 3 days after inoculation with 10^5^ PFU of the EV71 virus, EV71- 3D-E105NAEK, or Sinovic-EV71 vaccine. (**j, k**) Fluorescent images and H&E staining of small intestines of neonatal mice at day 14. Scale bars, 200 μm. Data in (**b–i**) are presented as mean ± SD (n = 5). Two-way analysis of variance (ANOVA) and Tukey’s test were performed. \*\*\*\**P* < 0.0001.

### *In vivo* UAA-controllable RNA virus evoked adjustable immune response in PylRS-tRNA^Pyl^ –transgenic mice

To investigate NAEK-controllable RNA viral replication *in vivo* with safeguards, we constructed transgenic mice stably harboring MmPylRS/tRNA^MmPyl^ via genome editing as previously described^19, 26–28^ to complement the *in vitro* UAA-controllable system (**Fig. 4a**). The expression of MmPylRS, tRNAMmPylCUA, and GFP^39TAG^ was confirmed in the small intestines of transgenic mice (**Fig. 4b, c, Extended Data Fig. 9**). Next, we administered daily i.p. injections to these mice of EV71-3D-E105NAEK and NAEK at different doses (−, no, 0 mg; +, low, 16 mg; ++, medium, 25 mg; +++, high, 50 mg) according to estimated bioavailability in the mice’s serum and small intestines (**Extended Data Fig. 10**). Three days later, we detected increased viral-RNA levels in an NAEK dose-dependent manner in the small intestine, brain, and spinal cord (**Fig. 4d**). Meanwhile, accumulating EV71-3D^P^°^l^ proteins were clearly observable by immunostaining and the pathological changes to the intestine shown by H&E staining **(Fig. 4e, f)**. No adult mice died from any NAEK or EV71-3D-E105NAEK dose; no obvious BW loss or other health issue in the low- and medium-dose groups was observed, as opposed to the high-dose group (**Fig. 4g, h**). Serum IgM and anti-VP1 IgG were robustly induced in a dose-dependent manner, even in the low-dose group (**Fig. 4i**), which was consistent with the boost dose of commercial Sinovic-EV71 vaccine (**Fig. 3e, f**), indicating the optimal dose for enhanced protective efficacy and few health concerns. These results demonstrated that the replication of EV71-3D-E105NAEK was NAEK controllable and dose-dependent in transgenic mice, further implying that EV71-3D-E105NAEK has great potential as a safe vaccine candidate. A single dose of EV71-3D-E105NAEK followed by optional UAA uptake could be the basis of a feasible immunization program to provoke optimal immune response in susceptible individuals with distinct levels of immunity, instead of injection of boost and even booster doses.

**Figure 4.**
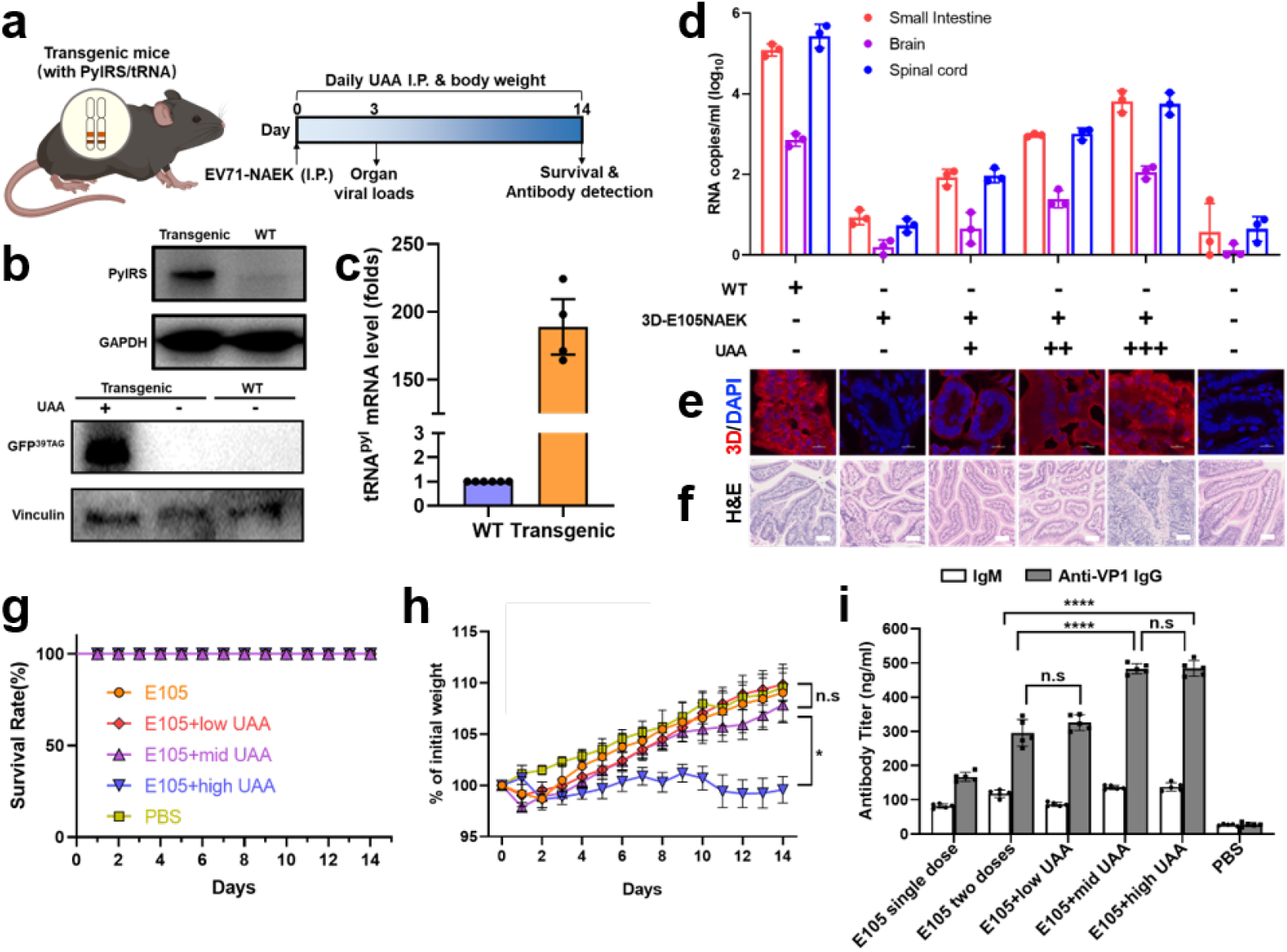
Immune response to UAA-controllable EV71-3D-E105NAEK in transgenic mice. (**a**) Transgenic mice harboring the UAA incorporation system were first injected i.p. with EV71-3D-E105NAEK for 14 days in the presence or absence of UAA. (**b**) WB analysis of PylRS and GFP^39TAG^ expression in transgenic mice. (**c**) Quantitative RT-PCR (qRT-PCR) analysis of tRNA^Pyl^ levels in transgenic mice. (**d**) Detection of viral-RNA copies in the small intestine, brain, and spinal cord after inoculation with 10^5^ PFU EV71-3D-E105NAEK in the presence or absence of UAA. (**e, f**) Fluorescent images and H&E staining of the small intestines of neonatal mice on day 14. Scale bars, 200 μm. (**g, h**) Survival rates and BW changes in transgenic mice after inoculation with EV71-3D-E105NAEK. (**i**) IgG and IgM titers detected after immunization with EV71-3D-E105NAEK in the presence of low-, medium-, or high- dose UAA, indicating in the UAA dose-dependent manner. Data (**d, g–i**) are presented as mean ± SD (n = 5). Two-way ANOVA and Tukey’s test were performed. \**P* < 0.05, \*\*\*\**P* < 0.0001.

## Discussion

The success of this viral-engineering strategy (1) defined guidelines for UAA incorporation and viral generation that could be broadly expanded to other RNA viruses in this study (2) provided a potential solution to the immediate quandary of vaccination by harnessing EV71-NAEK to elicit strong immune responses that were adjustable via artificial NAEK administration. For UAA-mediated viral engineering, although genetic codon expansion technology has been employed to modify virus-like particles^29, 30^, engineer viral vectors for gene therapy^31, 32^, and probe the biology of viruses^33, 34^, UAA- incorporated sites are generally selected by random mutations within targeted viral proteins, a traditional screening method for protein engineering^35^, and the lack of defined rules considering both protein modification and virus assembly. Moreover, engineered viruses developed in previous studies were always packaged in HEK293T cells with transiently or stably expressed PylRS/tRNA pairs^18^, which is appropriate for generation of common viral vectors (*e.g.*, adeno-associated viruses [AAVs], lentiviruses) but not for most viruses’ propagation, especially in vaccine development^36^.

Herein we reported that the established two-round viral-engineering system, consisting of NAEK-HEK293T cells for transfection and NAEK-Vero cells for propagation, could generate UAA-engineered viruses for main intestinal pathogens, including enteroviruses and coxsackieviruses, with high efficiency. Based on parallel screening in the EV-71 genome, we took chemical similarity, gene conservation, viral-protein property, and engineered viral titer into consideration when defining rules for UAA incorporation via logistic-regression analysis.

The final principle can be adapted to the diverse set of viable UAAs^37^ to generate engineered viruses with different modes of incorporation. Beyond the control of their replication, viral phenotypes can be simultaneously modified by the versatile side chain of UAA to jointly produce several desirable properties, such as improved cell tropisms and manufacturability^38, 39^. This is substantially more challenging than traditional methods of genetic engineering.

Additionally, UAA-engineered viruses restore the full infectious form for direct priming of the immune system, and they contain the full genome, which can recapitulate RNA replication and protein translation without virion assembly *in vivo*, like RNA-based vaccines in principle^40^. Therefore, we chose EV71-NAEK to validate that its safety and efficacy were comparable with those of currently available vaccines. In our mouse model, few viral RNA copies were detected even in the EV71-NAEK– vaccinated group. However, as maturing biotechnologies and extensive immunological discoveries advance the development of next-generation vaccines, the issue remains that policy-consistent vaccinations would be improper for individuals with poor immunity, especially children^41^, older adults^42^, and pregnant women^43^. The alternative solution is to choose a vaccine that might induce weak immune responses or change the dose and interval of vaccination; both strategies would fall short of the desired protective efficacy and would be impractical in large-scale applications.

Our study highlighted that the amendable EV71-NAEK could infect and replicate *in vivo* to elicit immune responses under control. To our knowledge, this is the first demonstration to date of vaccination with a controllable pathogen to evoke adjustable antibody landscapes. Compared with those of commercial vaccines, the absolute antibodies varied from 0.5-fold to approximately 2.5-fold in an external NAEK dose-dependent manner, suggesting that efficient protection can be achieved via an optimal dose of UAA according to the patient’s level of immunity. Currently, UAA-engineered virus retaining the full infectious form presents diverse antigens that, within artificial thresholds, would efficiently prime the human immune system with transient delivery of PylRS-tRNA^Pyl^ pairs and oral UAA uptake^19^, a novel and convenient vaccine strategy for combating emerging viruses. More generally, with advances in synthetic biology in smart materials and cell therapies^44^, the development of synthetic gene circuits for sensing immune responses and outputting UAAs to control viral replication could become the basis of a potential immunization program.

In conclusion, we systematically explored a novel strategy to convert infectious viruses with unnatural amino acids to achieve controllable viral replication and elicit an adjustable immune response *in vivo*, potentially creating a next-generation vaccine candidate. This study has laid the foundation for bioengineering design of smart vaccine candidates responsive to immunity, drawing on both synthetic biology and viral engineering.

## Materials and Methods

### Cell culture

The human embryonic kidney 293T (HEK293T, CRL-11268) cells and Vero-E6 (CRL- 1586) cells were obtained from ATCC. HEK293T cells were cultured in Dulbecco’s modified Eagle’s medium (DMEM) (Life Technologies) supplemented with 10% fetal bovine serum (FBS) (Gibco), 100 IU/mL penicillin and 100 μg/mL streptomycin (Invitrogen). Vero-E6 cells were cultured in Eagle’s Minimum Essential Medium (EMEM) (Life Technologies) with the same supplements. The cells were cultured at 37 °C and under 5% CO2, and passaged upon reaching 80%-90% confluence. Authentication and test for the free of mycoplasma were performed by our lab.

### Virus and vaccine

Enterovirus 71 AH/08/06 (Genbank: HQ611148.1, genotype C4) was used as a study model. The enterovirus type 71 vaccine (inactivated) and the EV71_SD059 strain lethal strain was provided by Sinovac Biotech (Beijing).

### Plasmid construction

The infectious full-length cDNA clone of EV71-A12 genome was kindly provided by the Academy of Military Medical Sciences. Mutant plasmids (EV71-A12-3C-TAG, EV71-A12-3D-TAG) containing amber codons within the open reading frame were generated by the QuikChange method (SBS Genetech) and confirmed by gene sequencing (Tsing ke, Beijing). The infectious full-length cDNA clone of CA10, CA16 and EVD68 were kindly provided by Xiamen University and were substituted with the same SP6-promoter transcription vector as EV71-A12. The Nucleocapsid protein sequence from SARS-CoV-2 (isolate Wuhan-Hu-1) was synthesized in BGI and fused to TagBFP gene by Gibson assembly. The *Methanosarcina mazei* (Mm) pyrrolysyl tRNA synthetase/tRNACUA pair (MmPylRS/tRNA^MmPylRS^) for site-specific incorporation of Nε-2-azidoethyloxycarbonyl-L-lysine (NAEK) was synthesized as previously reported^19^. The Tol2 transposon systems were purchased from Biocytogen (Beijing, China). The MmPylRS gene driven by CAG promoter was cloned into the Tol2 transposon vector to obtain Tol2-pylRS plasmid. The GFP gene with an amber codon introduced at residue position 39 was expressed under a CMV promoter and cloned into the Tol2-pylRS plasmid to obtain Tol2-pylRS-GFP^39TAG^ plasmid. The pcDNA 3.1 vectors carrying 6 tRNA^MmPylRS^ copies driven by human H1, human U6 and human 7sk promoters respectively were preserved in our lab. Then it was used as template to arrive the 12 tRNA^MmPylRS^ copies which were then cloned into the Tol2-pylRS-GFP^39TAG^ plasmid via Gibson assembly to obtain the Tol2-12tRNA^pyl^-pylRS-GFP^39TAG^ plasmid, used for NAEK-Vero stable cell line construction subsequently. All plasmids used for transfection were amplified using a Maxiprep kit (Promega) according to the manufacturer’s instructions.

### Establishment of NAEK-293T stable cell line

HEK293T cells were used for lentiviral vector packaging and transduction. The cells were cultured in DMEM medium (Macgene, without sodium pyruvate), supplemented with 10% FBS (PAA), and 1 mM nonessential amid acids (Gibco) in 6-well plates until sub-confluent. Then, cells were co-transfected with 0.72 µg of pSD31 transfer plasmid, 0.64 µg of pRSV, 0.32 µg of pMD2G-VSVG and 0.32 µg of pRRE using the transfection reagent Megatran1.0 (Origene). After 6 h, the transfection medium was replaced by DMEM medium containing 3% FBS and 1 mM nonessential amid acids. 48 h post-infection, the lentivirus-containing supernatant was harvested and filtered through a 0.45 µm filter. The resultant dual lentiviruses pSD31-PylRS and pSD31- GFP^39TAG^ were used to integrate MmPylRS and the GFP^39TAG^ gene into the genome of HEK293T cells. Experiments for stable lentiviral transduction were carried out as follows: HEK293T cells were seeded in a 6-well plate and transduced with lentiviral filtrates in the presence of 8 µg/mL polybrene 24 h later. Then, selection was performed under the pressure of 600 ng/mL puromycin and 200 µg/mL hygromycin until parental cells completely died. The resultant stably transduced HEK293T-PylRS/GFP^39TAG^ cells were transfected with linearized bjmu-12tRNA^MmPyl^ -zeo plasmid DNA and cultured under the pressure of 200 µg/mL Zeocin until parental cells completely died. In the presence of UAA, the stably transfected cells were then sorted by fluorescence-activated cell sorting (FACS) according to the GFP phenotype and verified by their dependence on UAA for GFP expression.

### Establishment of NAEK-Vero stable cell line

For electroporation, Vero-E6 cells were resuspended in Opti-MEM and counted, and 1×10^6^ cells were mixed with 15 μg of linearized Tol2-12tRNA^pyl^-pylRS-GFP^39TAG^ plasmid in medium cup (EC002S). The electroporation was performed using a CUY21EDIT square-wave electropulser (Nepa Gene Co., Ltd) under the following parameters: 150 V, poring pulse (5 ms with 50 ms intervals); 20 V, transfer pulse (50 ms with 50 intervals). After electroporation, the cells were immediately moved from the cup and plated on three 10-cm diameter tissue culture dishes in complete medium with 20% FBS. After 24 h, the plates were washed twice with Opti-MEM and re-fed with complete medium (10% FBS). To obtain stably transfected Vero, the cells were first incubated in complete medium with 20% FBS for 72 h and then in complete medium with 4 mM NAEK and 2 μg/ml puromycin. Nearly after 13 days, the Vero cells containing the transfected gene exhibited a phenotype to form clones. Then the pooled clones were cultured under puromycin pressure. In 15 days, about 80% of the Vero cells expressed GFP fluorescence, indicating MmPylRS/tRNA^MmPylRS^ gene was functional to read-through the amber codon introduced in GFP. The cells were sorted by FACS to obtain the monoclonal stable cell line.

### Generation of wild type EV71 viruses and UAA-controllable EV71 viruses

The wild type EV71 generated from reverse genetic system was performed as previous report^45^. 2x10^5^ cells per well from the HEK293T-tRNA/PylRS/GFP^39TAG^, HEK293T- 6GlnstRNA/GFP^39TAG^ and HEK293T-3CD/GFP cell lines were seeded into six well plates in DMEM supplemented with 10% FBS for 24 h before transfection. Then the wild type EV71 plasmid was linearized by MluI restriction enzyme digestion. The 1 μg linearized plasmid was serving as a template for *in vitro* transcription (SP6, Promega), and the products was then transfected into cells with Lipofectamine 2000 (Invitrogen). Six hours later, the medium containing the mixture of mRNA and lipofectamine 2000 reagent was replaced with DMEM supplemented with 1% FBS. The cells were further incubated at 37 °C in 5% CO2 until >90% CPE was observed, and the supernatant containing the generated virus was harvested and centrifuged at 1000 g for 10 min to remove contaminating cells.

To generate EV71-NAEK virus, an almost identical procedure was carried out, with the following changes: The plasmid expressing wild type viral RNA was replaced by the corresponding mutant plasmid, and the medium was further supplemented with 1 mM NAEK when the virus was packaged by UAA system for viral packaging. To identify the UAA dependent viral strains, a parallel packaging experiment was conducted without UAA supplement.

### Western blot analysis

HEK293T cells were lysed in RIPA lysis buffer (Applygen) supplemented with complete protease inhibitor cocktail (Roche) 48 h after transfection, and cell debris was removed by centrifugation. Tissue was then broken with a homogenizer and removed by centrifugation at 4 °C. Protein was extracted with lysis buffer and quantified by BCA assay (Thermo). A total of 100 µg protein from each sample was boiled with loading buffer, separated on 4%-12% NuPAGE (Invitrogen), and then electroblotted onto a polyvinylidenedifluoride membrane. The membrane was blocked with 5% (v/v) nonfat milk in TBST (50 mM Tris-HCl, 150 mM NaCl, and 0.02% Tween-20, pH 7.5) at room temperature for 1 h and then incubated with rabbit/mouse polyclonal antibodies overnight at 4 °C. Anti-3D antibodies (1:500, ab15277, Abcam), anti-N protein antibodies (1:500, ab7164, Abcam), anti-GFP antibodies (1:3000, 12715-1-AP, Proteintech), anti-PylRS antibodies (1:3000, sc-365062, Santa Cruz Biotechnology, to detect Myc-tagged PylRS) were diluted in TBST containing 5% (v/v) of defatted milk. The membranes were rinsed three times with TBST and then incubated with horseradish peroxidase-conjugated goat anti-rabbit/mouse IgG (1:3000) at room temperature for 1h. After extensive washing, the protein bands were developed using an enhanced chemiluminescent detection kit (Millipore). The optical bands were visualized in a Fuji Las-3000 dark box (FujiFilm), and band densities were quantified using Quantity One Analysis software. All western blot experiments were done for at least three times in parallel and the representative results were reported.

### Flow cytometry

Until transfected cells were treated with antibiotics to form clones, cells were dissociated into single cell using trypsin/EDTA and analyzed on BD FACSAriaTM (BD Biosciences) with the appropriate filter settings (488 nm coherent sapphire laser for GFP excitation). The front and side scatters were used to identify intact cells and mean background fluorescence from untransfected cells was subtracted from the measured signal. GFP positive cells were harvested for further validation of integrated genes.

### Virus growth curve analysis

To determine *in vitro* virus growth rates, triplicate wells of confluent transgenic cells, HEK293T-tRNA/PylRS/GFP^39TAG^, HEK293T-6stRNA^Gln^/GFP^39TAG^ and HEK293T-3CD/GFP, (6-well plate format, 10^6^ cells/well) were infected at a MOI of 0.1. After 1 h of virus adsorption at 37 °C, cells were washed and overlaid with DMEM supplemented with 1% FBS, and additional treatment with 1 mM NAEK in UAA system. At the indicated times post-infection (on day 1, 2, 3, 4, 5, 6 and 7), the cell supernatants were collected and viral titers were determined by the qRT-PCR as described above.

### Enterovirus 71 quantification by 50% tissue culture infective dose

For quantifying all viruses’ stocks, the 50% tissue culture infective dose (TCID50/mL) titers were determined. In brief, 5×10^4^ Vero-E6 cells (or NAEK-Vero cells for EV71- NAEK quantification) were seeded in 96-well plates the day before infection. The virus samples were serially diluted with DMEM containing 1% FBS (10^3^ to 10^10^) and then each of dilution was added in wells separately. The plates were incubated at 37 °C in 5% CO2 for 2-5 days. The cytopathic effect (CPE) was observed under a microscope and virus titer was determined using the Reed-Münch endpoint calculation method (http://www.fao.org/3/AC802E/ac802e0w.htm).

### Generation of MmPylRS/tRNA^MmPyl^ transgenic mice

Methods to generation of transgenic mice are previously described^19, 27, 28^. In brief, we chose the ROSA26 site of the autosome and utilized *Streptococcus pyogenes* Cas9 (SpCas9) and designed sgRNA to cut the targeted genome, which was then recombined with the transgene of CAG pro-MmPylRS-IRES-EGFP^39TAG^-WPRE-PA-(7SK-tRNA ^MmPyl^) 4 in the pronucleus of fertilized eggs derived from parental mice (C57BL/6♂ x C57BL/6♀). Subsequently, we performed embryo transplantation into surrogate ICR females to breed the F0 generation. Three weeks after birth, the tail DNA of all the F0 generation mice was extracted for PCR analysis to identify the positive mice, which were further crossed with wild type C57BL/6 mice (♀) to produce the next generation.

### Animal experiments

All animal experiments were performed in accordance with the guidelines of the Institutional Animal Care and Use Committee of the Peking University. Briefly, groups of 4-week old female BALB/c mice (n=5) were inocubated by intraperitoneal injection with EV71-NAEK virus, Sinovic EV71 vaccine (positive control) and PBS (negative control) respectively to test the immunogenicity, efficacy and safety. All Mice were monitored daily for body weight and survival rate. Two weeks post-inoculation, five mice of each group were sacrificed. The sera were used to test their immunogenicity by ELISA and organs (e.g., brain, intestine and skeletal muscle) were collected to detect viral titer by qRT-PCR.

For protective efficacy study, five BALB/c mice of 4-week old in each group were immunized for twice (0 and 14 days) with the same dose. Two weeks after the second immunization, the female mice were paired with male mice until the born of neonatal mice. Four-day old neonatal mice were intraperitoneally injected with 10 LD50 of EV71_SD059 strain (lethal strain, Sinovac Biotech, Ltd, Beijing). The body weight and survival rate of mice were observed for 7-14 days until the mice returned to normal.

For *in vivo* UAA-controllable infection investigation, eight transgenic mice of 4-week old in each group were intraperitoneally injected with EV71-3D-E105NAEK, wild type EV71 virus and PBS after genotype identification by PCR analysis. All mice were intraperitoneally injected with different doses of NAEK and monitored daily for body weight and survival rate. Three days after infection, three mice of each group were sacrificed to detect viral titer by qRT-PCR and pathological changes by immunofluorescence staining and histological analysis. Other five mice of each group were sacrificed in two weeks to collect their sera for neutralizing antibodies detection.

### Quantitative Real-Time PCR analysis

Total RNA harvested from cells or mice tissues infected with distinct viruses or vaccine was extracted by using Trizol reagent (Invitrogen, CA, USA). RNA (1 μg) was subjected to RT-PCR in accordance with the protocol provided by Promega. The transcripts were quantitated and normalized to the internal GAPDH control. The primers used in the experiments are listed in **Extended Data Table 3**. The PCR conditions were 1 cycle at 95 °C for 5 min, followed by 40 cycles at 95 °C for 15 s, 60 °C for 1 min, and 1 cycle at 95 °C for 15 s, 60 °C for 15 s, 95 °C for 15 s. The results were calculated using the 2^-△△CT^ method according to the GoTaq qPCR Master Mix (Promega) manufacturer’s specifications.

### The whole transcriptome analysis

RNA sequencing was performed by Annoroad Inc. (Beijing, China). Briefly, total mRNA was purified by poly-T oligo-attached magnetic beads and fragmented. Sequencing libraries were generated using NEBNext® Ultra™ RNA Library Prep Kit for Illumina® (#E7530L, NEB, USA) following the manufacturer’s recommendations and index codes were added. The libraries were sequenced on an Illumina platform and 150 bp paired-end reads were generated. The raw data were quality controlled by Q30 and the clean data was aligned to the reference genome (Ensembl hg38) using HISAT2 (v2.1.0) software. Reads Count for each gene in each sample was counted by HTSeq v0.6.0, and FPKM (Fragments Per Kilobase Millon Mapped Reads) was then calculated to estimate the expression level of genes in each sample. DESeq was used for differential gene expression analysis between two samples and Genes with q≤0.05 and |log2_ratio| ≥ 1 were identified as differentially expressed genes. GO and KEGG pathway enrichment analysis were performed based on differentially expressed mRNAs.

### Enzyme-linked immunosorbent assay (ELISA)

IgM and anti-VP1 IgG antibody in sera were measured using ELISA^46^. In this assay, 96-well ELISA plates (Thermo Fisher Scientific Inc., USA) were coated with 100 mM bicarbonate/carbonate buffer (pH 9.6) containing 0.03 μg/mL of recombinant proteins (VP1) from homologous wild-type viruses (Sino Biological Inc., Beijing, China) overnight at 4 °C, followed by 3% bovine serum albumin (BSA; Sigma) in PBS-0.05% Tween 20 (blocking buffer) blocking for 1 h at 37 °C. The coated plates were then incubated with serum samples for IgM and anti-VP1 IgG, and followed by incubating HRP-conjugated anti-mouse/ferret IgG antibody or HRP-conjugated anti-mouse/ferret IgM antibody (Sino Biological Inc., Beijing, China) for 1 h at 37 °C. Plates were detected with tetramethyl benzidine (TMB) substrate (Millipore, Billerica, MA, USA) and stopped after 15 min with 0.5 M H2SO4. Optical density (O.D.) was read at 450 nm using an ELISA reader (Microplate Reader AMR-100; Allsheng, Hangzhou, China).

### Immunofluorescence

The intestine harvested from different mice were fixed in 4% PFA at room temperature for 1 hour, followed by immersing in 30% (w/v) sucrose until submersion before embedding and freezing in the Optimal Cutting Temperature (OCT) compound (Tissue-Tek). Serial 12 µm sections were obtained by cryo-sectioning of the embedded intestine tissue at -20 °C using a cryostat (Leica). Cryosections were blocked with 5% normal donkey serum (Jackson ImmunoResearch) in PBST for 30 min. The sections were incubated with anti-3D antibody (1:2000, GTX630193, GeneTex) diluted in blocking buffer at 4 °C overnight. The slides were subsequently incubated with secondary goat anti-rabbit IgG Alexa Fluor 594 (1:400, LifeTech) at room temperature for 1 h. The slides were stained with 0.5 µg/mL Hoechst and mounted in mounting media. Stained sections were photographed under a Nikon Ti-S microscope, and the dystrophin-positive cells were analyzed.

### Histological analysis

The tissues of different mice were isolated and fixed in 4% PFA solution for two days, sequentially dehydrated with 70%, 95% and 100% ethanol, and defatted with xylene for 2 h before being embedded in paraffin. The 10-μm-thick section was cut and subjected to hematoxylin and eosin staining (H&E staining). The slices were observed under optical microscopy, and the histological morphology of different mouse groups was compared.

### Statistical analysis

Data are presented as mean ± SD. In most circumstances, three replicates were used for each group unless otherwise noted. The statistical significance between two groups was determined by student’s t test. For three or more groups, one-way or two-way ANOVA with Tukey’s multiple comparisons test was performed. Homogeneity of variances among groups was confirmed using Bartlett’s test. Conformity to normal distribution was confirmed using the Kolmogorov-Smirnov test. All tests were two-sided and performed in GraphPad Prism 8. A probability of p < 0.05 was considered as statistically significant. For annotations, * p<0.05; ** p<0.01; *** p<0.001; **** p<0.0001. The amino acid similarity was calculated in Maestro of the Schrodinger software. Logistic regression was performed between the virus packaging results and incorporated positions features.

## Reporting summary

Further information on research design is available in the Nature Research Reporting Summary linked to this paper.

## Data availability

The authors declare that all data supporting the results in this study are available within the paper and its **Extended data figure legends**.

## Acknowledgments

We thank all the lab members’ contribution to this work. Q. X. was supported by the National Science and Technology Major Projects for “Major New Drugs Innovation and Development” (No. 2018ZX09711003-001-003, 2018-2020).

## Author contributions

Q. X. conceived the idea and designed the experiments. Z. Z, Y. W. and X. W. performed the experiments and organized the results, Z. Z., X.W., X. Y., H. L, H. Z., and Q. X. analyzed the data, prepared the figures and wrote the manuscript. All other authors participated in some of the experiments, results or discussion. Q. X. supervised the study. Z. Z., Y. W., X. W. and H. L. contributed equally to this work.

## Additional information

**Supplementary information** is available for this paper at https://doi.org.

**Correspondence and requests for materials** should be addressed to Q. X.

## Extended Data Figures

**Extended Data Figure 1.**
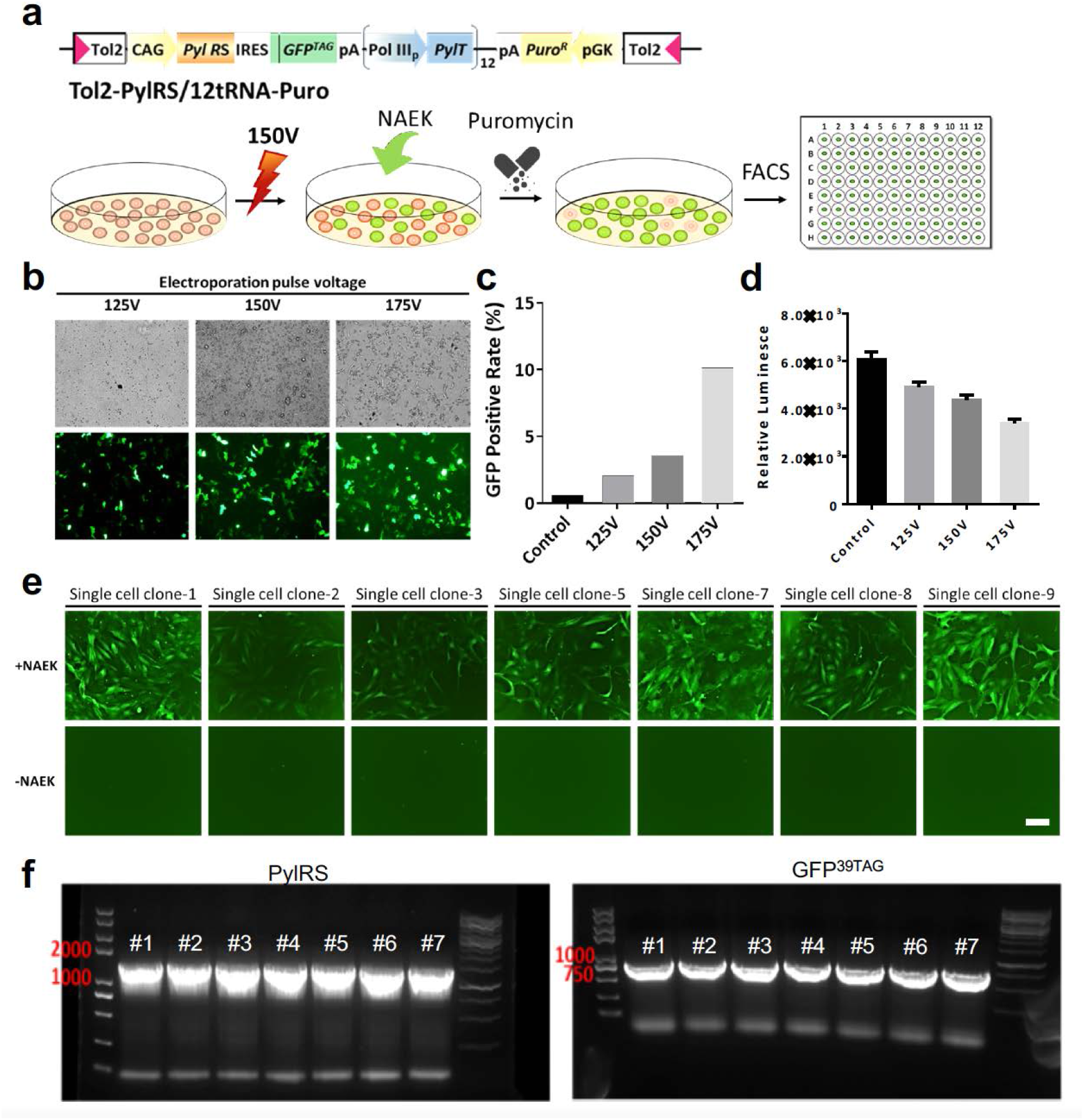
Establishment of the VeroE6-tRNA^pyl^/PylRS/GFP^39TAG^ stable cell line. (**a**) Plasmid of the UAA incorporation system and stable cell line screening. (**b**) Comparison of transfection efficiency at different voltages. (**c, d**) Number of green fluorescent protein (GFP)–positive cells and rate of relative luminescence at different voltages. (**e**) GFP fluorescence appeared in the screened stable cell line in a NAEK-dependent manner. Scale bars, 50 μm. (**f**) PCR analysis of PylRS and GFP expression in the stable cell line.

**Extended Data Figure 2.**
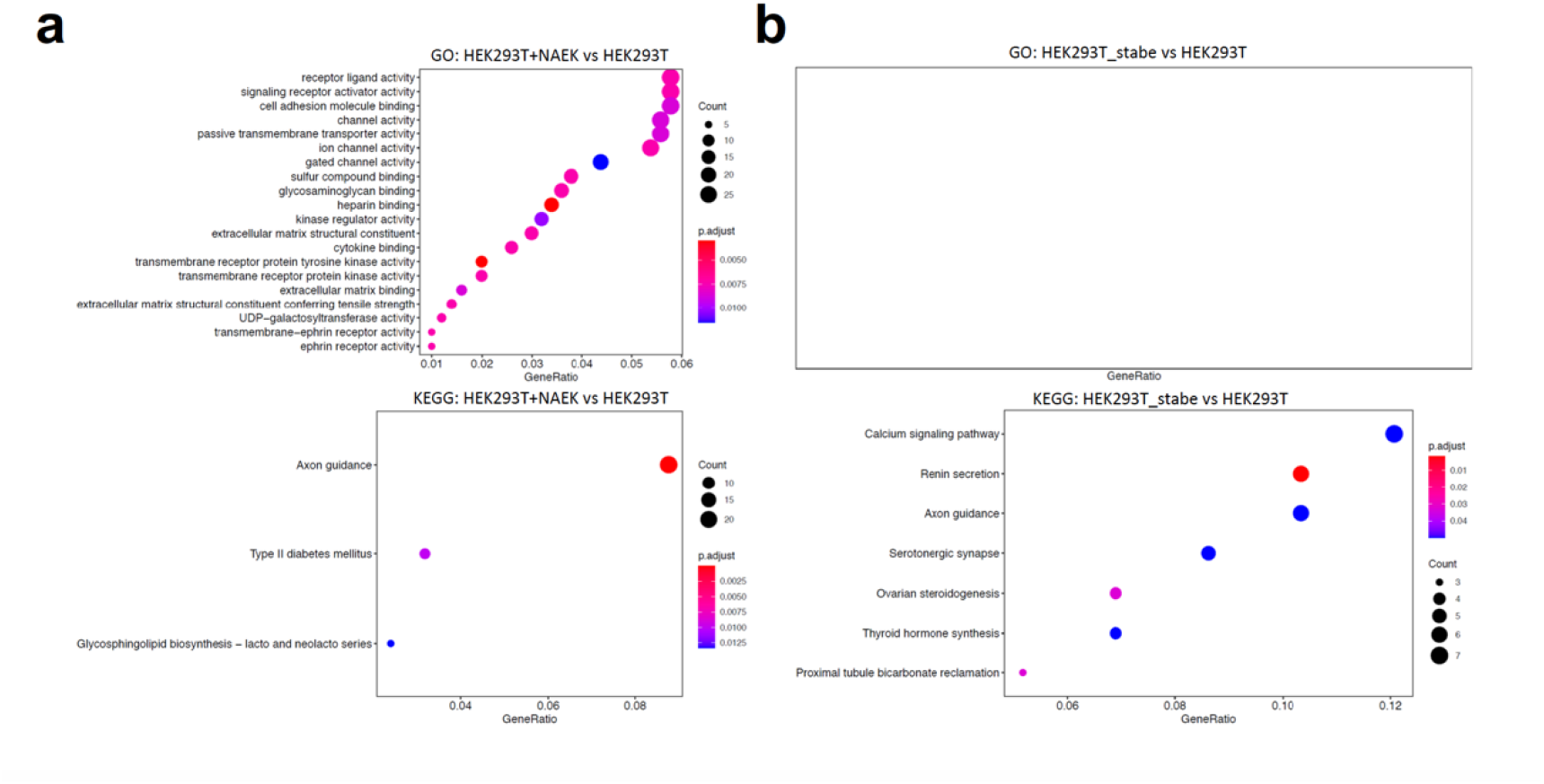
Representative GO terms and KEGG enrichment analysis of HEK293T+NAEK cells and the engineered packaging systems compared with HEK293T cells.

**Extended Data Figure 3.**
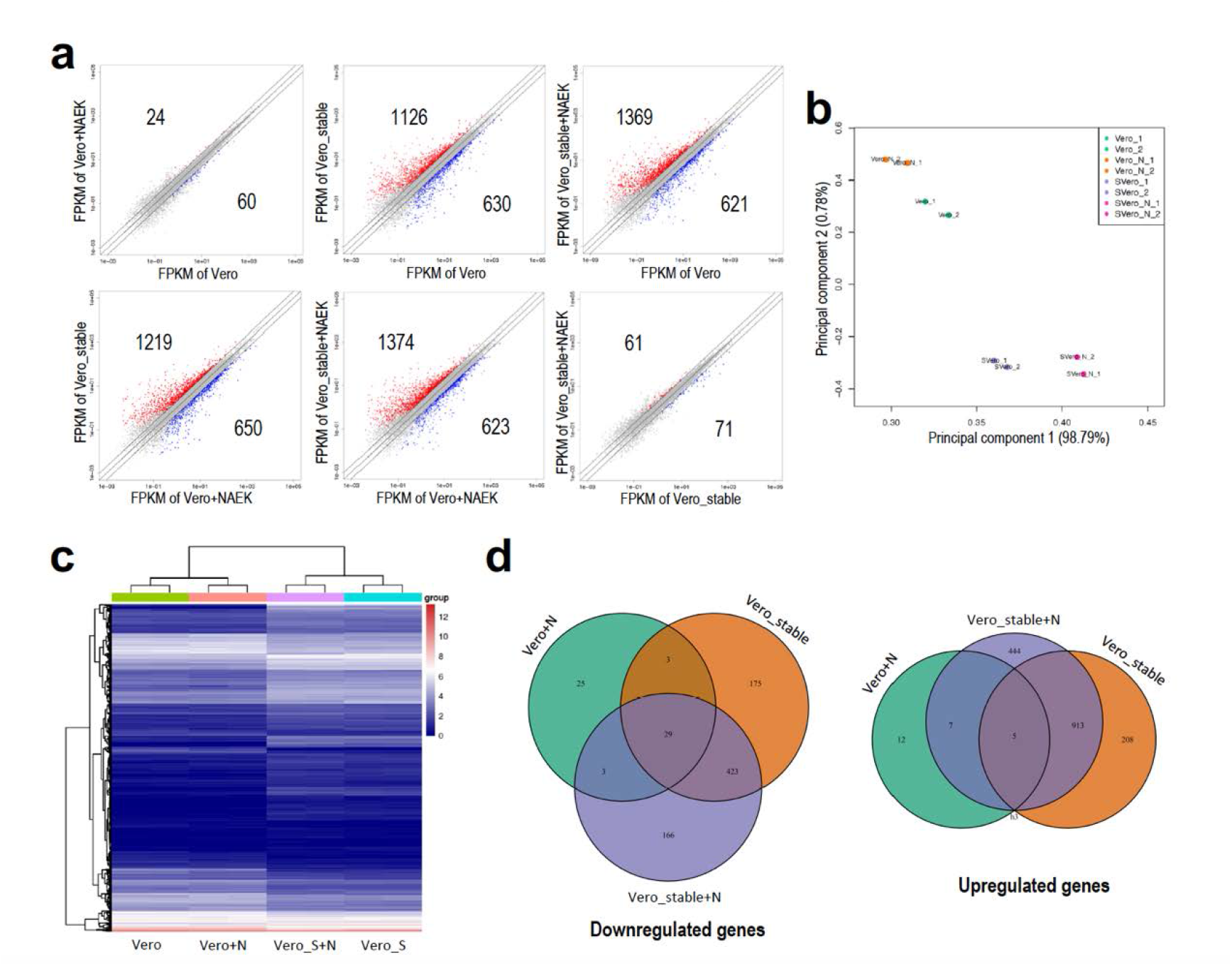
Whole-transcriptome analysis of Vero cells and the engineered production systems. (**a**) Plots show whole-transcriptome FPKMs. Red dots indicate upregulated genes; blue dots downregulated genes. Two biological replicates were used per sample. (**b**) PCA of Vero cells and the engineered production systems. (**c**) Hierarchical-clustering and heatmap analysis of DEGs in Vero cells and the engineered production systems. (**d**) Venn diagram showing significant overlap (*P* < 0.005) of DEGs between Vero cells and the engineered production systems.

**Extended Data Figure 4.**
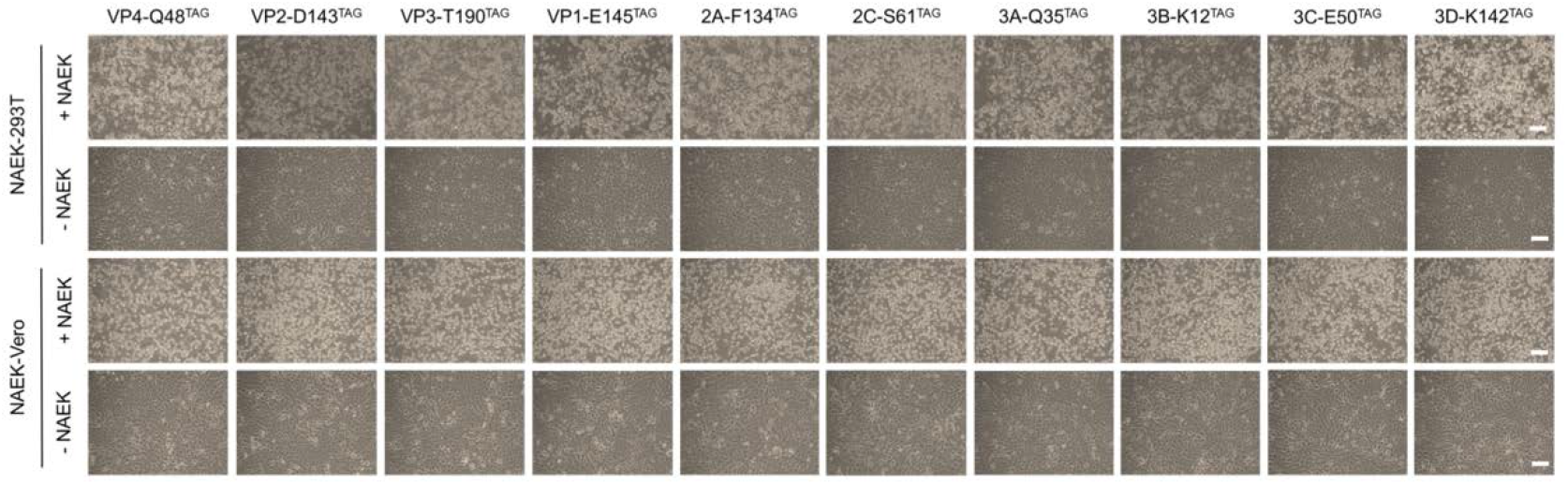
CPE of EV71-NAEK virus harboring an amber codon at VP1–VP4, 2A–2C, and 3A–3D by random-site selection. Scale bars, 50 μm.

**Extended Data Figure 5.**
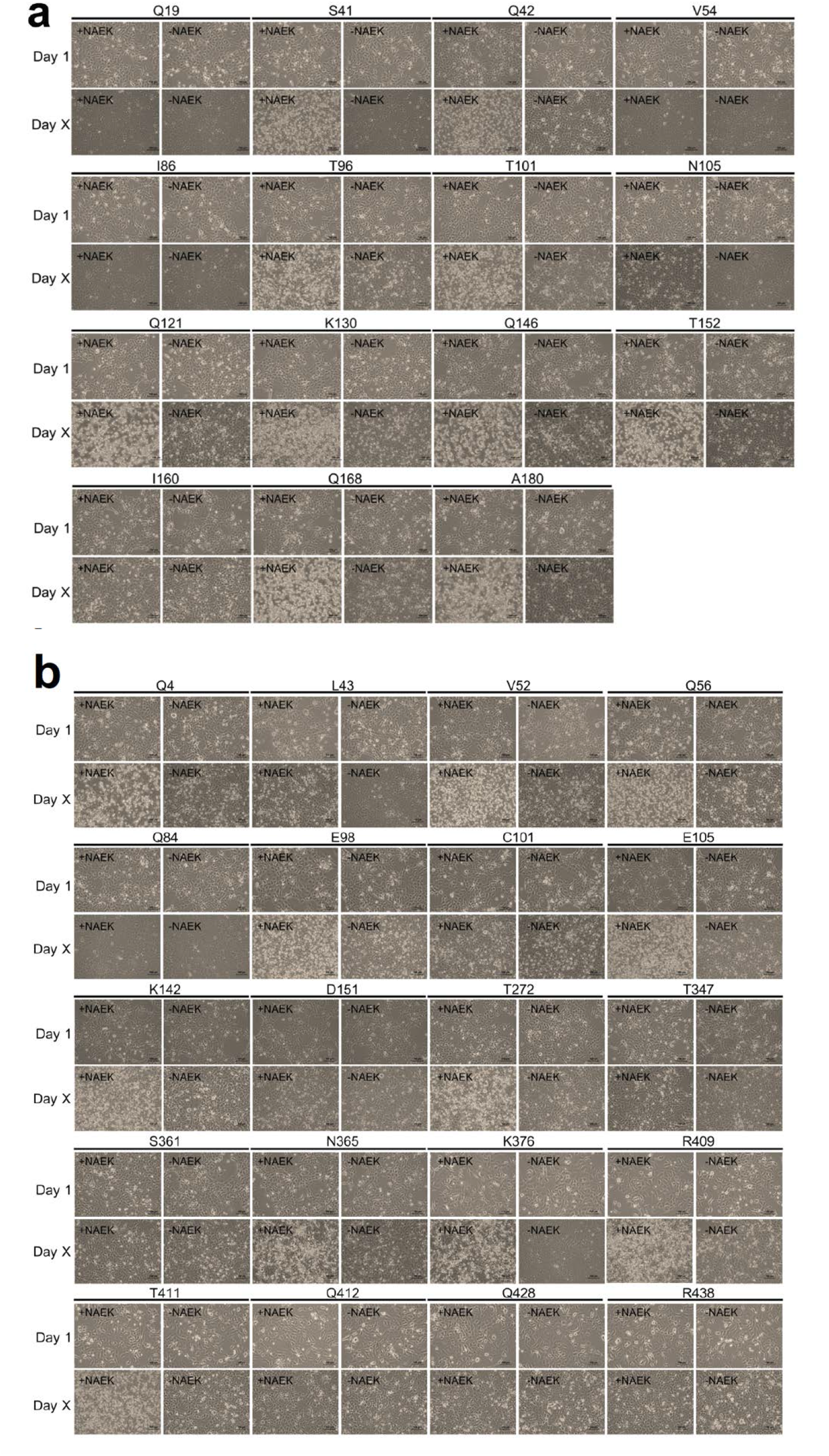
CPE imaging of EV71-NAEK strains produced by the NAEK incorporation system in the presence or absence of NAEK. Day X represents days required to attain ∼100% CPE (n = 3). (**a**) Strains with NAEK incorporated into 3C protein. (**b**) Strains with NAEK incorporated into 3D protein.

**Extended Data Figure 6.**
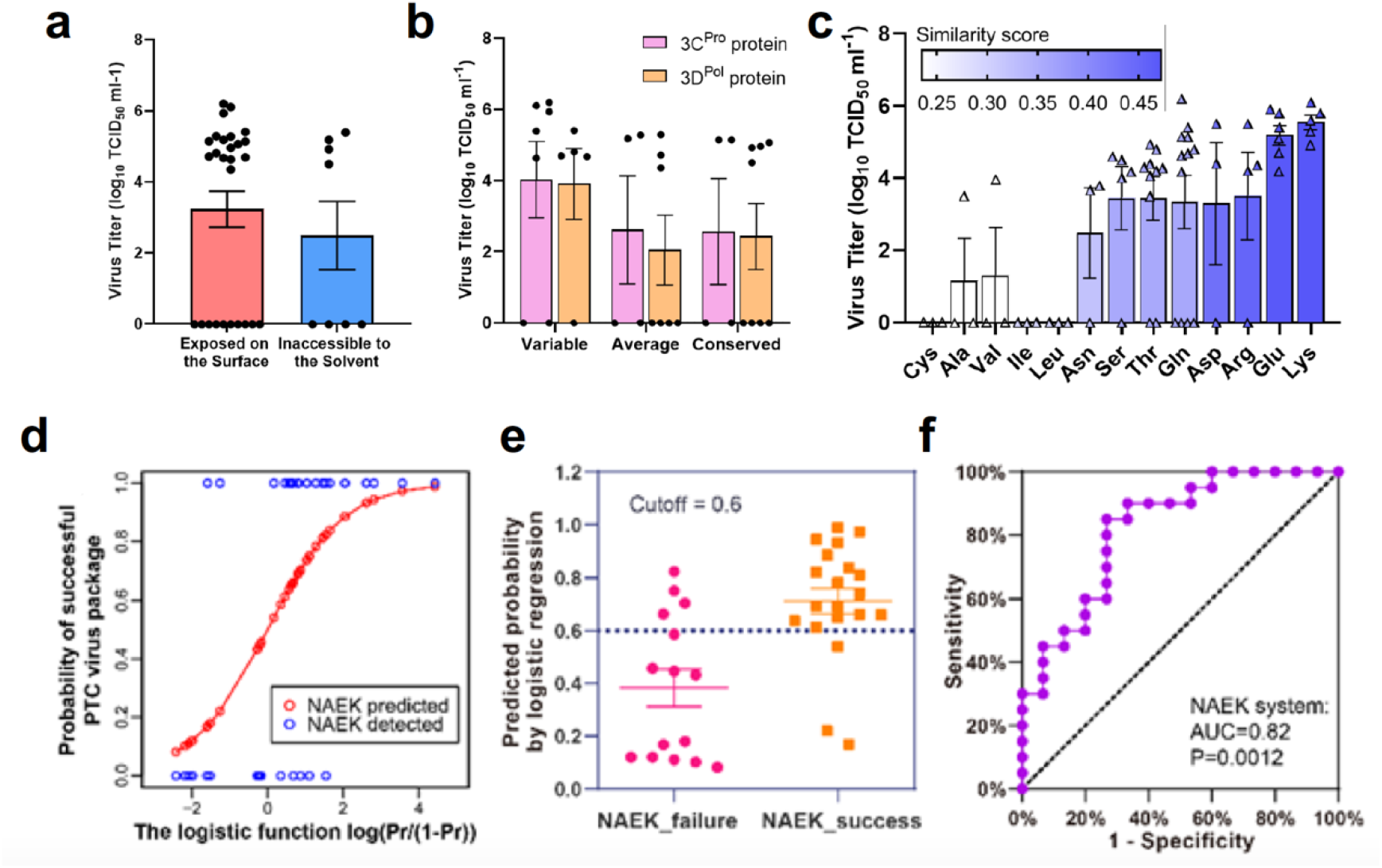
Guideline for and validation of NAEK-controllable virus design for other single-stranded RNA (ssRNA) viruses. (**a–c**) Relationship between viral titers and residue location, and conservation or similarity of amino acids. (**d**) Workflow for the guideline for and validation of controllable virus design. (**e–g**) Logistic regression of viral packaging by NAEK, based on the information of the 35 environmentally verified positions in EV71. Blue circles indicate the detected probability (0 or 1) of successful viral packaging, red circles the predicted probability thereof. Four factors were selected for logistic regression for function and related estimates. Predicted probability of success or failure per group was also plotted (mean ± standard error of the mean [SEM]), and receiver operating characteristic (ROC) curves are shown (area under the curve [AUC] > 0.8).

**Extended Data Figure 7.**
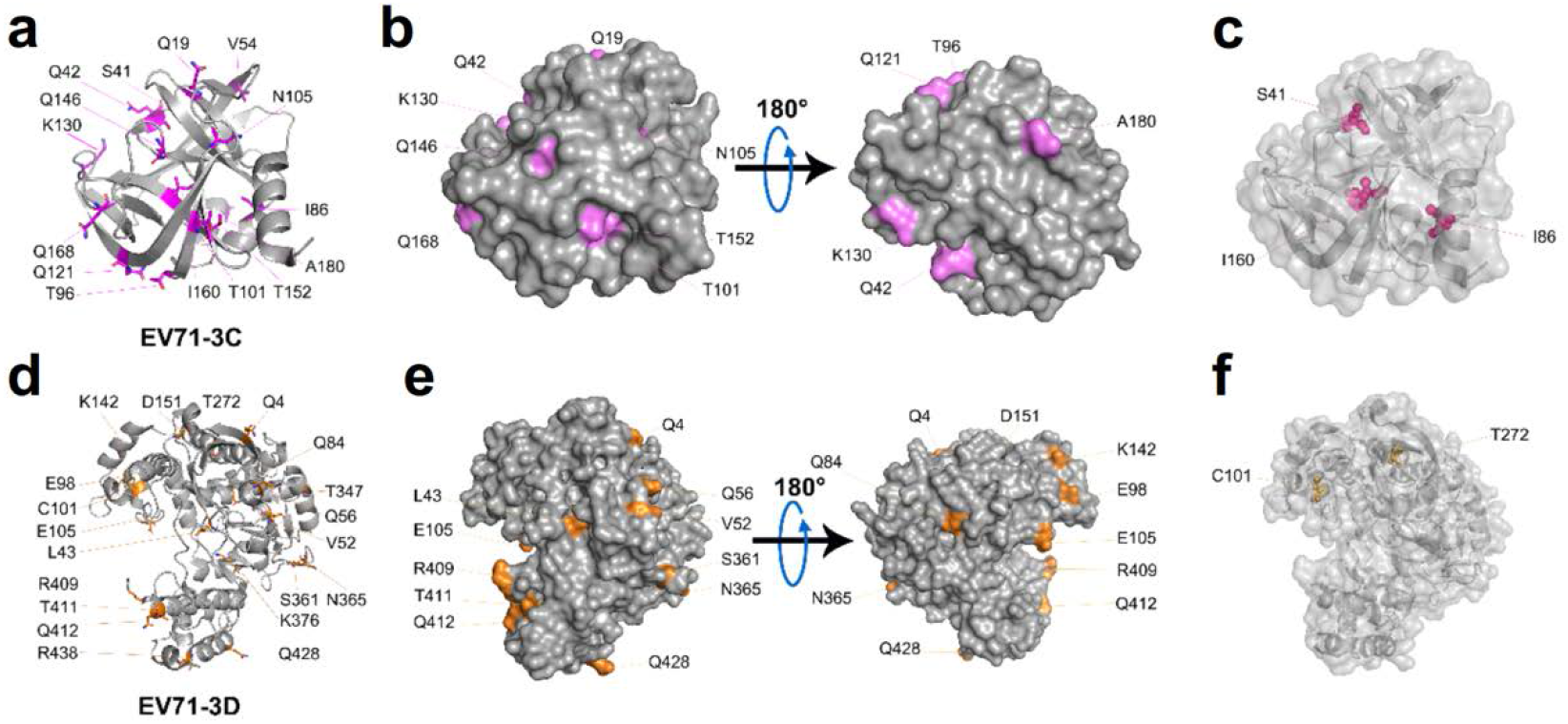
Structural illustration of 35 residues on EV71 3C/D. (**a**) Residue location on EV71 3C protein (PDB: 4GHQ). (**b**) Residues on 3C surface are highlighted in magenta. (**c**) Residues located within 3C are shown in the sphere. (**d**) Residue location on EV71 3D protein (PDB: 3N6L). (**e**) Residues on 3D surface are highlighted in orange. (**f**) Residues located within 3D are shown in the sphere.

**Extended Data Figure 8.**
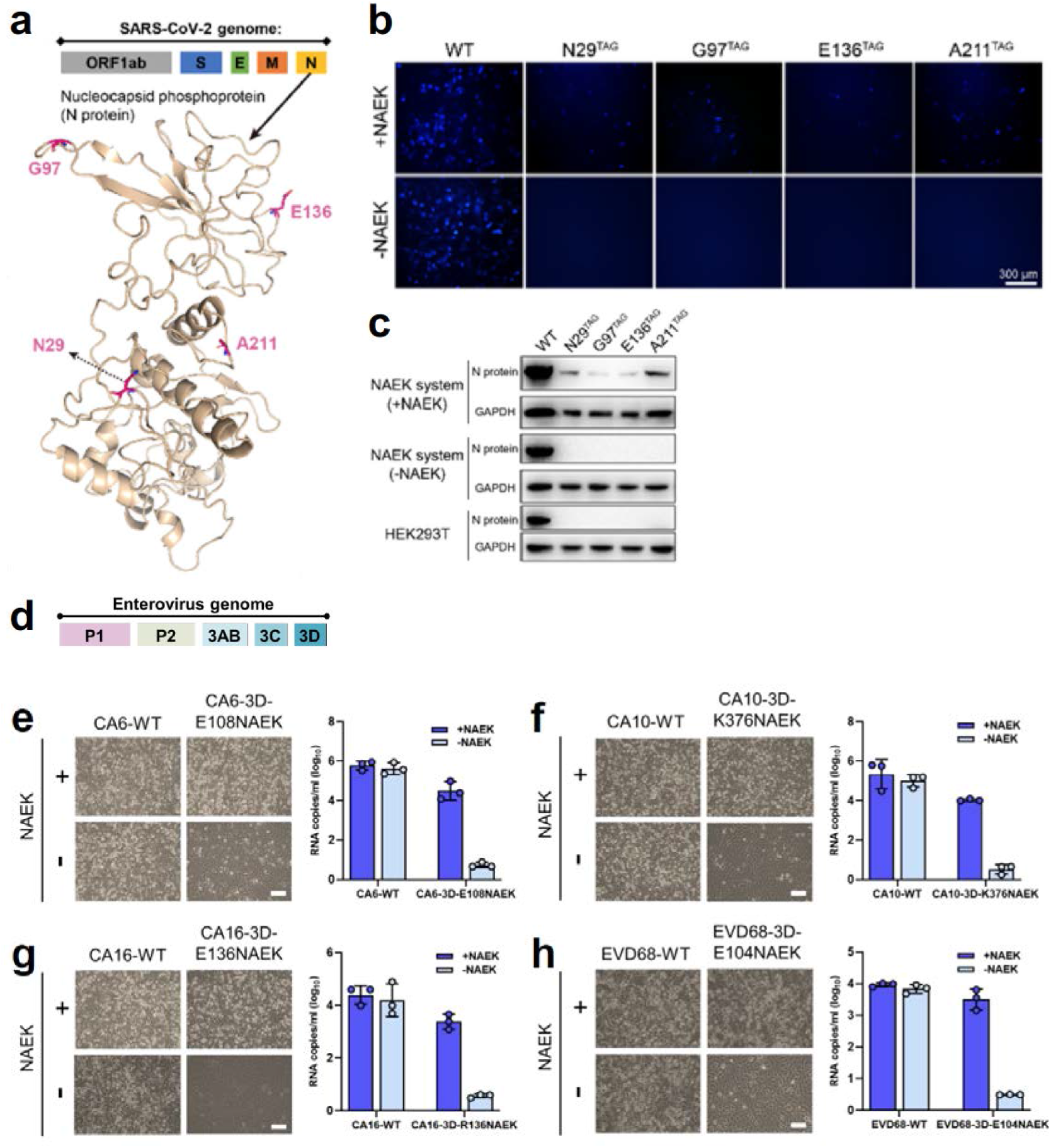
(**a**) Schematic of SARS-CoV-2 protein arrangement and structural model of four target residues on the N protein. The gold pentagram indicates the location of N in the viral genome. The N protein structure was modeled using the Phyre2 online server (Kelley LA et al., *Nature Protocols* 10, 845–858 [2015]). (**b**) Fluorescent imaging of expression of the four recombinant N proteins. (**c**) WB analysis of expression of the four recombinant N proteins. (**d**) Structure of the *Enterovirus* genome. (**e**) UAA-dependent CPE formation and RNA copies of CA6-3D-E108NAEK, CA10-3D-K376NAEK, CA16-3D-E136NAEK, and EVD68-3D-E104NAEK.

**Extended Data Figure 9.**
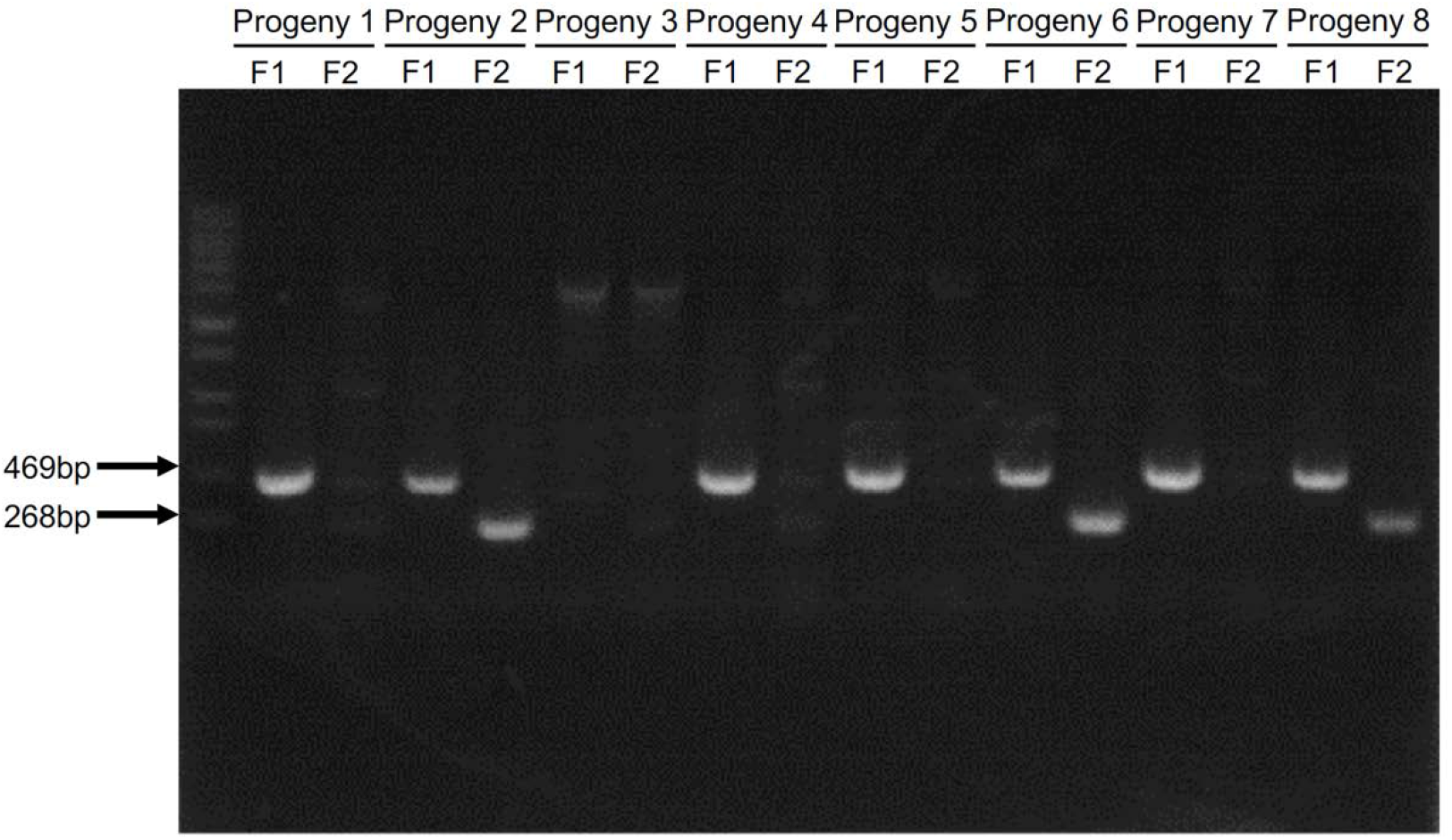
PCR analysis of mouse tail DNA from six descendants of transgenic mice crossed with wild-type mice. We used two primers; Primer 1 targeted the sequence at 900 bp, Primer 2 at 300 bp. The six descendants were from the F1 generation. Results showed that descendants 5 and 6 were positive for effective PylRS-tRNA^Pyl^ pairs integration.

**Extended Data Figure 10.**
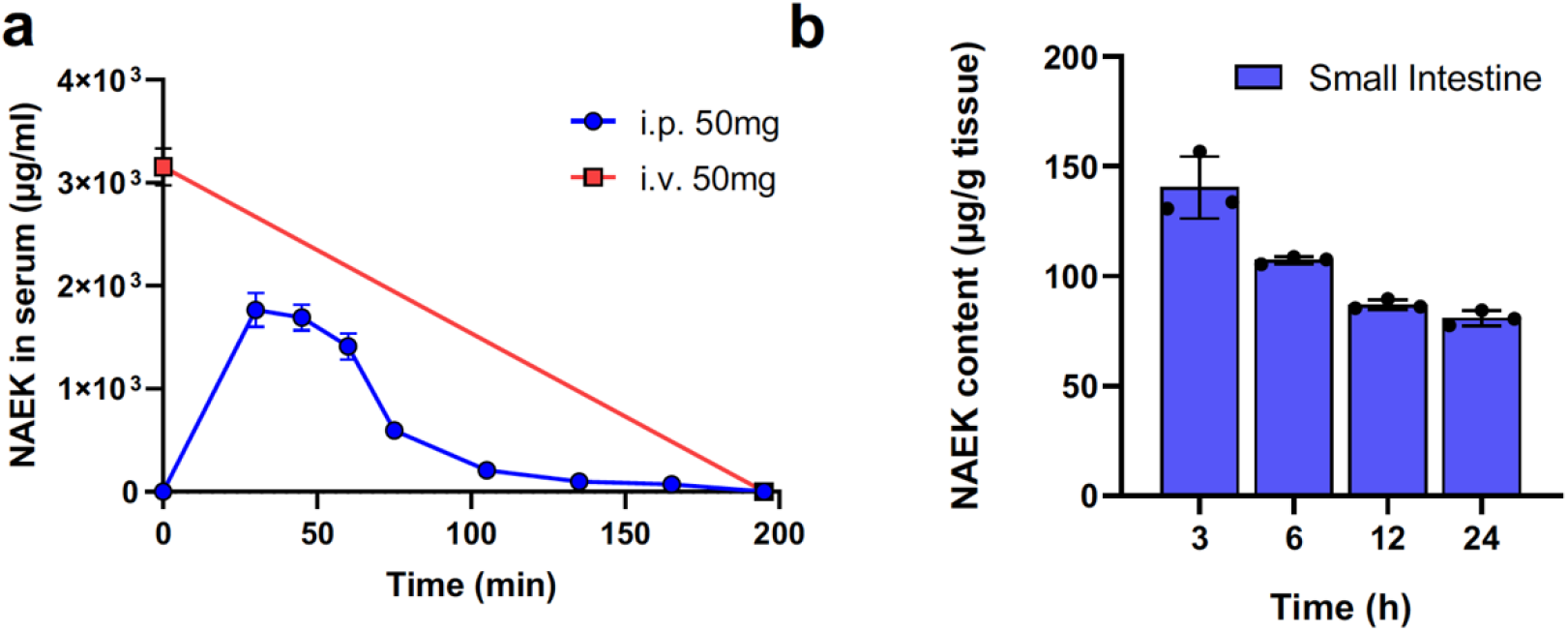
Bioavailability of NAEK. Bioavailability of NAEK (administered i.p. and i.v.) in (**a**) serum and (**b**) tissue (small intestine) was as estimated by AUC. One 500-mg dose of NAEK was administered. Data are presented as mean ± SD (n = 3).

## Reference

1. Baltimore, D. Expression of animal virus genomes. Bacteriol. Rev. 35, 235–241 (1971).

2. Koff, W. C. et al. Accelerating Next-Generation Vaccine Development for Global Disease Prevention. Science 340, 1232910 (2013).

3. Thanh Le, T., et al. The COVID-19 vaccine development landscape. Nat. Rev. Drug Discov. 19, 305–306 (2020).

4. Lurie, N., Saville, M., Hatchett, R. & Halton, J. Developing Covid-19 Vaccines at Pandemic Speed. N. Engl. J. Med. 382, 1969–1973 (2020).

5. Ketzer, P., et al. Artificial riboswitches for gene expression and replication control of DNA and RNA viruses. Proc. Natl. Acad. Sci. 111, E554–E562 (2014).

6. Chung, H. K. et al. Tunable and reversible drug control of protein production via a self-excising degron. Nat. Chem. Biol. 11, 713–720 (2015).

7. Tahara, M., et al. Photocontrollable mononegaviruses. Proc. Natl. Acad. Sci. 116, 11587–11589 (2019).

8. Heilmann, E. et al. Chemogenetic ON and OFF switches for RNA virus replication. Nat. Commun. 12, 1362 (2021).

9. Jeyanathan, M. et al. Immunological considerations for COVID-19 vaccine strategies. Nat. Rev. Immunol. 20, 615–632 (2020).

10. Excler, J.-L., Saville, M., Berkley, S. & Kim, J. H. Vaccine development for emerging infectious diseases. Nat. Med. 27, 591–600 (2021).

11. Thi Nhu Thao, T., et al. Rapid reconstruction of SARS-CoV-2 using a synthetic genomics platform. Nature 582, 561–565 (2020).

12. Xie, X. et al. An Infectious cDNA Clone of SARS-CoV-2. Cell Host Microbe 27, 841–848.e3 (2020).

13. Arunachalam, P. S. et al. Systems vaccinology of the BNT162b2 mRNA vaccine in humans. Nature 596, 410–416 (2021).

14. Cho, A. et al. Anti-SARS-CoV-2 receptor binding domain antibody evolution after mRNA vaccination. Nature 1–9 (2021) doi:10.1038/s41586-021-04060-7.

15. Chin, J. W. et al. An Expanded Eukaryotic Genetic Code. Science 301, 964–967 (2003).

16. Wang, L., Brock, A., Herberich, B. & Schultz, P. G. Expanding the Genetic Code of Escherichia coli. Science 292, 498–500 (2001).

17. Wang, N., et al. Construction of a Live-Attenuated HIV-1 Vaccine through Genetic Code Expansion. Angew. Chem. Int. Ed. 53, 4867–4871 (2014).

18. Si, L. et al. Generation of influenza A viruses as live but replication-incompetent virus vaccines. Science 354, 1170–1173 (2016).

19. Shi, N. et al. Restoration of dystrophin expression in mice by suppressing a nonsense mutation through the incorporation of unnatural amino acids. *Nat*. Biomed. Eng. 1–12 (2021) doi:10.1038/s41551-021-00774-1.

20. Chen, Y. et al. Controlling the Replication of a Genomically Recoded HIV-1 with a Functional Quadruplet Codon in Mammalian Cells. ACS Synth. Biol. 7, 1612– 1617 (2018).

21. Zhang, X. et al. A trans-complementation system for SARS-CoV-2 recapitulates authentic viral replication without virulence. Cell 184, 2229–2238.e13 (2021).

22. Ricardo-Lax, I. et al. Replication and single-cycle delivery of SARS-CoV-2 replicons. Science eabj8430 (2021) doi:10.1126/science.abj8430.

23. Pollard, A. J. & Bijker, E. M. A guide to vaccinology: from basic principles to new developments. Nat. Rev. Immunol. 21, 83–100 (2021).

24. Ooi, M. H., Wong, S. C., Lewthwaite, P., Cardosa, M. J. & Solomon, T. Clinical features, diagnosis, and management of enterovirus 71. Lancet Neurol. 9, 1097– 1105 (2010).

25. Solomon, T. et al. Virology, epidemiology, pathogenesis, and control of enterovirus 71. Lancet Infect. Dis. 10, 778–790 (2010).

26. Brown, W., Liu, J. & Deiters, A. Genetic Code Expansion in Animals. ACS Chem. Biol. 13, 2375–2386 (2018).

27. Chin, J. W. Expanding and reprogramming the genetic code of cells and animals. Annu. Rev. Biochem. 83, 379–408 (2014).

28. Chen, Y. et al. Heritable expansion of the genetic code in mouse and zebrafish. Cell Res. 27, 294–297 (2017).

29. ElSohly, A. M. et al. ortho-Methoxyphenols as Convenient Oxidative Bioconjugation Reagents with Application to Site-Selective Heterobifunctional Cross-Linkers. J. Am. Chem. Soc. 139, 3767–3773 (2017).

30. Behrens, C. R. et al. Rapid Chemoselective Bioconjugation through Oxidative Coupling of Anilines and Aminophenols. J. Am. Chem. Soc. 133, 16398–16401 (2011).

31. Erickson, S. B., et al. Precise Photoremovable Perturbation of a Virus–Host Interaction. Angew. Chem. Int. Ed. 56, 4234–4237 (2017).

32. Zheng, Y. et al. Broadening the versatility of lentiviral vectors as a tool in nucleic acid research via genetic code expansion. Nucleic Acids Res. 43, e73–e73 (2015).

33. Sakin, V. et al. A Versatile Tool for Live-Cell Imaging and Super-Resolution Nanoscopy Studies of HIV-1 Env Distribution and Mobility. Cell Chem. Biol. 24, 635–645.e5 (2017).

34. Nikić, I., et al. Minimal Tags for Rapid Dual-Color Live-Cell Labeling and Super- Resolution Microscopy. Angew. Chem. Int. Ed. 53, 2245–2249 (2014).

35. Wang, Y., et al. Directed Evolution: Methodologies and Applications. Chem. Rev. (2021) doi:10.1021/acs.chemrev.1c00260.

36. Kiesslich, S. & Kamen, A. A. Vero cell upstream bioprocess development for the production of viral vectors and vaccines. Biotechnol. Adv. 44, 107608 (2020).

37. Young, D. D. & Schultz, P. G. Playing with the Molecules of Life. ACS Chem. Biol. 13, 854–870 (2018).

38. Kelemen, R. E., et al. A Precise Chemical Strategy To Alter the Receptor Specificity of the Adeno-Associated Virus. Angew. Chem. Int. Ed. 55, 10645–10649 (2016).

39. Yao, T. et al. Site-Specific PEGylated Adeno-Associated Viruses with Increased Serum Stability and Reduced Immunogenicity. Molecules 22, 1155 (2017).

40. Chaudhary, N., Weissman, D. & Whitehead, K. A. mRNA vaccines for infectious diseases: principles, delivery and clinical translation. Nat. Rev. Drug Discov. 1–22 (2021) doi:10.1038/s41573-021-00283-5.

41. Lewnard, J. A., Lo, N. C., Arinaminpathy, N., Frost, I. & Laxminarayan, R. Childhood vaccines and antibiotic use in low- and middle-income countries. Nature 581, 94–99 (2020).

42. Mulligan, M. J. et al. Phase I/II study of COVID-19 RNA vaccine BNT162b1 in adults. Nature 586, 589–593 (2020).

43. Dagan, N. et al. Effectiveness of the BNT162b2 mRNA COVID-19 vaccine in pregnancy. Nat. Med. 27, 1693–1695 (2021).

44. Cubillos-Ruiz, A. et al. Engineering living therapeutics with synthetic biology. Nat. Rev. Drug Discov. 1–20 (2021) doi:10.1038/s41573-021-00285-3.

45. Roehl, H. H., Parsley, T. B., Ho, T. V & Semler, B. L. Processing of a cellular polypeptide by 3CD proteinase is required for poliovirus ribonucleoprotein complex formation. J. Virol. 71, 578–585 (1997).

46. Zhang, Z. et al. Immunogenicity and Safety of an Inactivated Enterovirus 71 Vaccine Administered Simultaneously With Hepatitis B Vaccine and Group A Meningococcal Polysaccharide Vaccine: A Phase 4, Open-Label, Single-Center, Randomized, Noninferiority Trial. J. Infect. Dis. 220, 392–399 (2019).

